# Isoxanthopterin: An Optically Functional Biogenic Crystal in the Eyes of Decapod Crustaceans

**DOI:** 10.1101/240366

**Authors:** Benjamin A. Palmer, Anna Hirsch, Vlad Brumfeld, Eliahu D. Aflalo, Iddo Pinkas, Amir Sagi, S. Rozenne, Dan Oron, Leslie Leiserowitz, Leeor Kronik, Steve Weiner, Lia Addadi

**Affiliations:** Department of Structural Biology, Weizmann Institute of Science, Rehovot, 7610001, Israel.; Department of Materials and Interfaces, Weizmann Institute of Science, Rehovot, 7610001, Israel.; Department of Chemical Research Support, Weizmann Institute of Science, Rehovot, 7610001, Israel.; Department of Life Sciences, Ben-Gurion University of the Negev, Beer-Sheva, 8410501, Israel.; The National Institute for Biotechnology in the Negev, Ben-Gurion University of the Negev, Beer-Sheva, 8410501, Israel.; Department of Organic Chemistry, Weizmann Institute of Science, Rehovot, 7610001, Israel.; Department of Physics of Complex Systems, Weizmann Institute of Science, Rehovot, 7610001, Israel.

## Abstract

The eyes of some aquatic animals form images through reflective optics. Shrimp, lobsters, crayfish and prawns possess reflecting superposition compound eyes, composed of thousands of square-faceted eye-units (ommatidia). Mirrors in the upper part of the eye (the distal mirror) reflect light collected from many ommatidia onto the underlying photosensitive elements of the retina, the rhabdoms. A second reflector, the tapetum, underlying the retina, back-scatters dispersed light onto the rhabdoms. Using microCT and cryo-SEM imaging accompanied by *in situ* micro-X-ray diffraction and micro-Raman spectroscopy, we investigated the hierarchical organization and materials properties of the reflective systems at high resolution and under close to physiological conditions. We show that the distal mirror consists of three or four layers of sparse plate-like nano-crystals. The tapetum is a diffuse reflector composed of hollow nanoparticles constructed from concentric lamellae of crystals. Isoxanthopterin, a pteridine analog of guanine, forms both the reflectors in the distal mirror and in the tapetum. The crystal structure of isoxanthopterin was determined from crystal structure prediction calculations and verified by comparison with experimental X-ray diffraction. The extended hydrogen bonded layers of the molecules results in an extremely high calculated refractive index in the H-bonded plane, *n* = 1.96, which makes isoxanthopterin crystals an ideal reflecting material. The crystal structure of isoxanthopterin, together with a detailed knowledge of the reflector superstructures, provide a rationalization of the reflective optics of the crustacean eye.

**Significance:** Aquatic animals use reflectors in their eyes either to form images or to increase photon capture. Guanine is the most widespread molecular component of these reflectors. Here we show that crystals of isoxanthopterin, a pteridine analogue of guanine, form both the image-forming ‘distal’ mirror and the intensity-enhancing tapetum reflector in the compound eyes of some decapod crustaceans. The crystal structure of isoxanthopterin was determined, providing an explanation for why these crystals are so well suited for efficient reflection. Pteridines were previously known only as pigments and our discovery raises the question of which other organic molecules may be used to form crystals with superior reflective properties either in organisms or in artificial optical devices.

## Introduction

Many spectacular optical phenomena exhibited by animals are produced by light-reflection from organic (1–3) or inorganic crystals (4, 5). A fascinating manifestation of such biological reflectors is in vision (6). Certain animals use mirrors instead of lenses to form images. This strategy is particularly useful in aquatic environments where the reduced refractive index contrast in water makes conventional lens-based eyes less effective. Furthermore, mirror-containing eyes are often extremely efficient light-collectors and are usually found in nocturnal animals or animals inhabiting dim-light environments (7). Two types of image-forming reflective eyes are found in nature: concave mirrored eyes and reflection superposition compound eyes. The former is epitomized by the scallop eyes, which produce well-resolved images by reflecting light from a concave, guanine crystal-based mirror located at the back of the eye onto the retina above it (8, 9). Similar eyes are also found in deep-sea fish (10, 11) and crustaceans (12, 13). The latter type of reflective eye, the reflecting superposition compound eye, is the focus of this study.

The reflecting superposition compound eye is found in decapod crustaceans (14, 15). Each compound eye is composed of thousands of square-faceted eye-units called ommatidia. An image is formed when light is reflected from the top of each ommatidium onto the retina, comprising the photo-sensitive ‘rhabdoms’ (Fig. 1) (16). In contrast to apposition compound eyes, where each ommatidium acts as an isolated optical unit, in the reflecting compound eye, light collected from many ommatidia is reflected across a ‘clear zone’ and is brought to focus as a single upright image on the convex-shaped retina (Fig.1C). The superposition of light rays from multiple eye units, effectively increases the pupil size making this an extremely light-sensitive device, which is well adapted for dim-light habitats. Reflection of light from the ommatidia operates in two regimes: grazing incidence light, with angles of incidence < ~15° relative to the ommatidial axis, is internally reflected from the walls of the ommatidia (15, 17). This is made possible by the refractive index contrast in the upper parts of the ommatidium (*n*=1.41 inside and *n*=1.34 outside the ommatidium in the region of the ‘crystalline cone’, Fig.1C) (15, 17). At larger angles of incidence, this index contrast is too small to lead to significant reflectivity. Light impinging at higher angles is thus reflected by a high-refractive-index square mirror (‘distal mirror’, Fig.1C), which surrounds each eye unit. Vogt postulated (14, 18) that the distal mirror of crustacean eyes is a multilayer reflector, but using conventional electron microscopy methods, he was unable to retain the *in vivo* structure. The square arrangement of the mirror acts like a corner reflector, whereby light incident from oblique planes will be reflected from two orthogonal mirrors by a total of 180°, and will return parallel to its original direction and be brought to focus on the retina (15, 17). A second reflector, the tapetum, lies immediately behind the retina and is responsible for the observed eye-shine of decapod crustaceans (19, 20). The tapetum reflects light back through the retina, giving the retina a second chance of absorbing light that was not absorbed on the first pass (6).

**Fig. 1.**
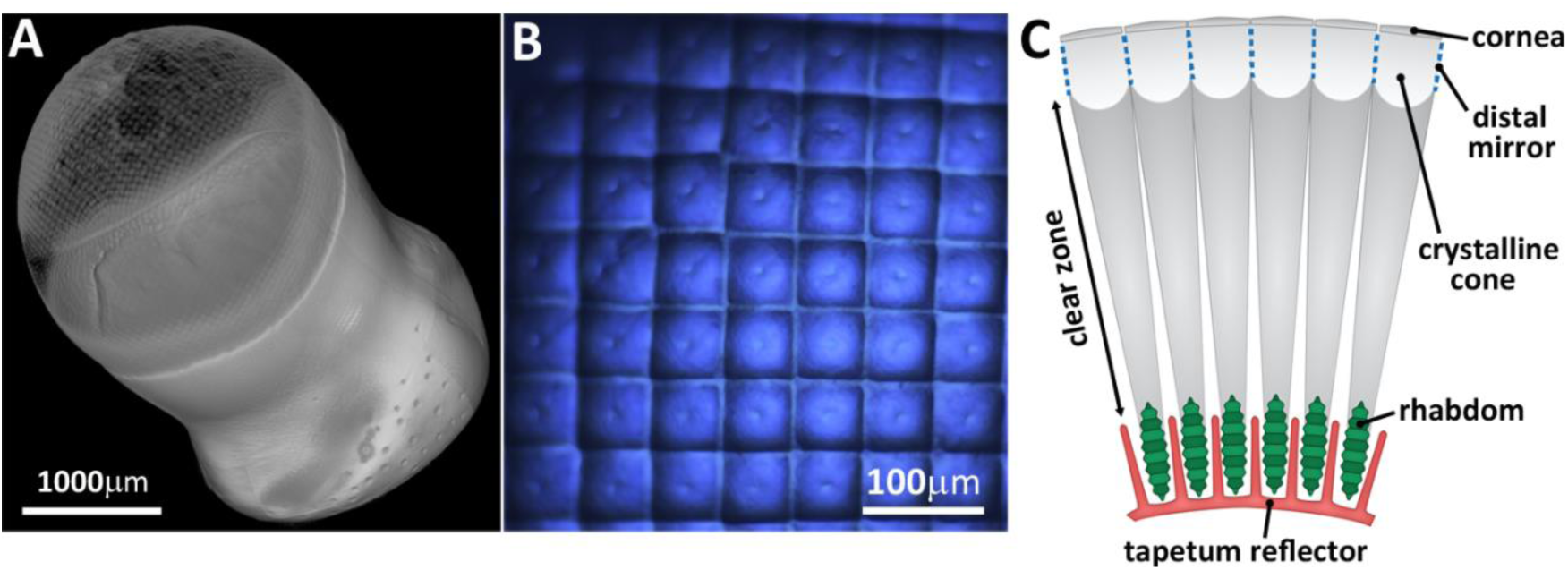
The reflecting superposition compound eye. (*A*) Volume rendering of an X-ray microCT scan of a whole crayfish eye (*Procambarus clarkii*). (*B*) Light microscopy image of the cornea looking down the eye axis from above. The characteristic square facets of the cornea (16) are clearly visible. (C) Schematic of the compound eye viewed perpendicular to the eye axis. Light transmitted through the cornea, is reflected off the distal mirror which lines the top parts of each eye unit (ommatidium) in the region of the crystalline cone (a homogeneous jelly-like material). The reflected light from many ommatidia passes across the clear zone and is brought to focus on the photoreceptor body of the retina – the rhabdom. A tapetum reflector lies behind the retina and contains projections which envelope the rhabdoms.

Surprisingly, little is known about the nature of the reflective materials in crustacean eyes. Using chromatographic and histochemical methods Kleinholz (21) identified xanthine, uric acid, hypoxanthine, xanthopterin and an unidentified pteridine in the retinal region of the lobster *Homarus americanus*. Using similar approaches Zyznar and Nicol (22) found isoxanthopterin in relatively high abundance in the vicinity of the distal and tapetum reflectors in the eyes of the white shrimp *Penaeus setiferus*. However, these studies could not determine the precise locations and phase of the purine and pteridine molecules. These uncertainties motivated our study of the properties of the reflective materials in the reflecting superposition eye.

We report that the reflectors in these eyes are composed of crystals of isoxanthopterin. We show that the distal mirror is a specular reflector formed from a few layers of plate-like nano-crystals. The tapetum is a diffuse reflector composed of hollow nanoparticles constructed from concentric lamellae of crystals which back-scatter light to the retina. By solving the crystal structure of isoxanthopterin, together with a detailed analysis of the hierarchical organization of the reflectors, we provide a rationalization for the reflective optics in this unique visual system.

## Results

We first determined the 3D organization of the eye components on the micro-to millimeter scale, using X-ray microCT measurements on fixed, dark-adapted eyes from a freshwater crayfish, *Cherax quadricarinatus* (Fig. 2A, B) and a freshwater prawn, *Machrobrachium rosenbergii*. Similar observations were made on both eyes, and are shown only for *C. quadricarinatus* in Fig. 2A, B. Two regions of high X-ray attenuation were identified in the upper part of the eye: the cornea and the distal mirror. The 45 *μ*m thick cornea, the most highly attenuating structure in the eye, forms a smooth boundary around the eye. The high density might well be due to the presence of some form of calcium carbonate (23). The cornea is formed from a mosaic of weakly-refracting square micro-lenses (24), about 50 *μ*m wide, with each micro-lens being associated with a single ommatidium (Fig. 1B). The second highly-attenuating feature is observed 60–65 *μ*m below the base of the cornea. When viewed perpendicular to the eye axis (Fig. 2A), this highly attenuating feature appears as a series of lines extending down the sides of each ommatidium in the region of the crystalline cone (Fig. 1C). When viewed down the eye-axis (Fig. 2B), the structure appears as an ordered array of squares (53 *μ*m wide), lining the square ommatidium. Polarized light-microscopy (Fig. 2C) reveals a highly birefringent material, about 100 *μ*m long, lining the crystalline cone, in the same region as the high-contrast structure observed in the X-ray microCT (Fig. 2B). The intensity of the birefringence varies significantly when the ommatidia are rotated with respect to cross-polarizers demonstrating that the material is composed of a highly oriented and most likely crystalline material (Fig. S1). The optical axes of the crystals are oriented along the ommatidial axis (17, 18).

**Fig. 2.**
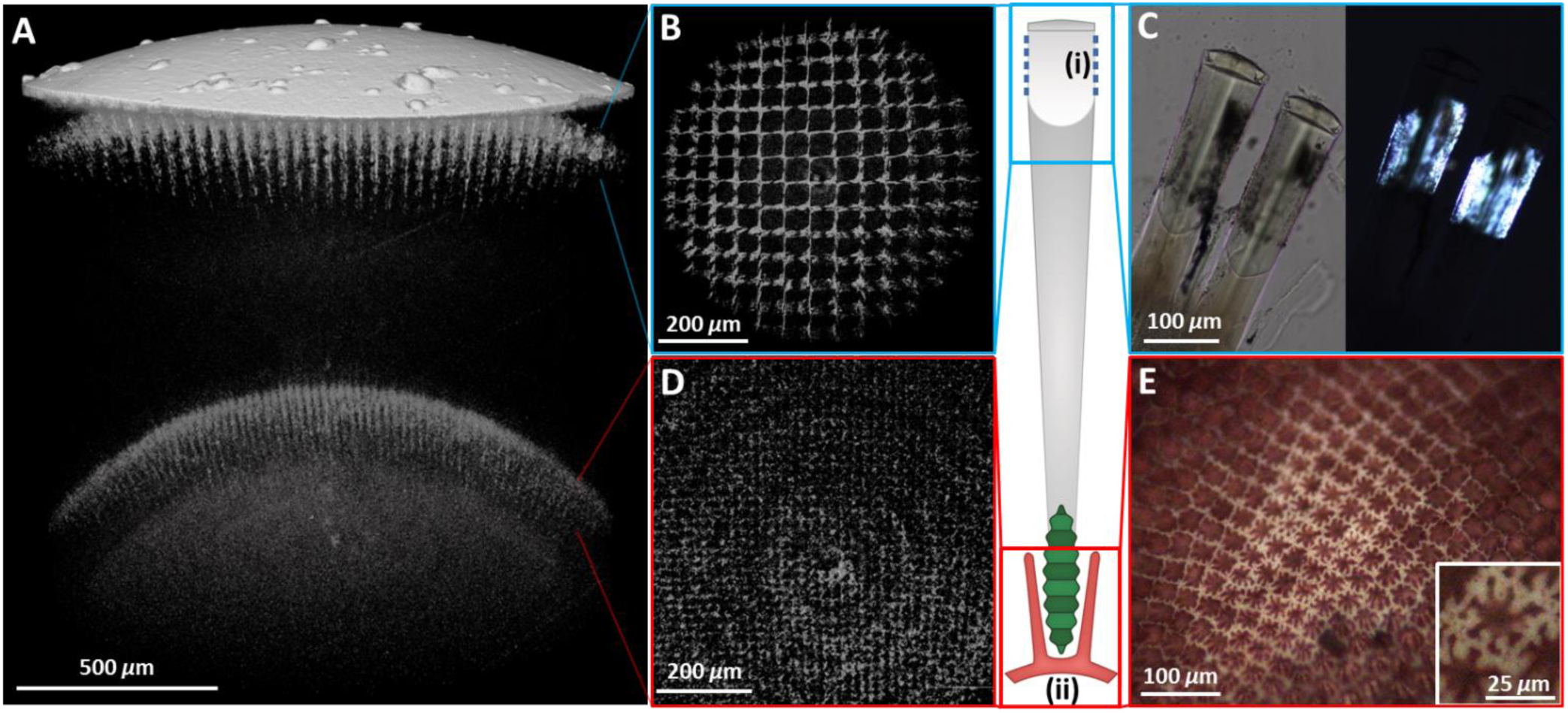
Micro- to millimeter scale architecture of the distal and tapetum reflectors. (*A*) X-ray microCT scan of a crayfish eye (*C. quadricarinatus*) viewed perpendicular to the eye axis, showing three high contrast features: the cornea (uppermost), distal mirror (below cornea) and tapetum (lower). (*B*) X-ray microCT image of the multilayer reflector underneath the cornea viewed from above, along the eye axis [blue square, schematic (i)]. (*C*) Light microscopy images of the upper parts of two ommatidia viewed perpendicular to the ommatidial axis [blue square, schematic (i)] without polarization (left) and between crossed-polarizers (right) showing a birefringent material lining the crystalline cone. (*D*) X-ray microCT image of the tapetum [red square, schematic (ii)] viewed from above and along the eye axis. (*E*) Tapetum viewed along the eye axis under polarized light [red square of schematic (ii)], showing the organization of the birefringent material. Inset: one of the square-prism structures within which the rhabdom and retinal cells reside. Schematic: A single ommatidium viewed perpendicular to the eye axis with the distal mirror (i) and tapetum (ii).

A region of low contrast (the ‘clear zone’ in Fig. 1C) separates the distal mirror from a third area of high X-ray contrast in the lower part of the eye, the tapetum (Fig. 2A, 1C). When viewed along the eye axis in microCT (Fig. 2D) and by light microscopy (Fig. 2E) the tapetum appears as a series of open squares *ca*. 25 *μ*m wide, containing a white birefringent material (Fig. 2E). The birefringence intensity does not vary upon rotation between crossed-polarizers indicating that the material is most likely crystalline, but the crystallites have no preferential orientation. The tapetal cells form a close sheath around the seven retinal cells and the seven-lobed rhabdom which is located in the center of the cavity (inset, Fig. 2E)(25–27). The high X-ray contrast observed in the microCT in the tapetum originates specifically from the reflective materials rather than with absorbing pigments residing above and below the tapetum (Fig. S2).

To ascertain if the tapetum material of *C. quadricarinatus* and *M. rosenbergii* is crystalline we performed *in situ* X-ray diffraction experiments using a microspot beam at the BESSY synchrotron. X-ray diffraction patterns obtained from the region of the rhabdoms (green spot, Fig.3A) exhibit no Bragg scattering, but do display a weak anisotropic scattering pattern at small angles consistent with the orthogonal ordering of the microvilli in the rhabdoms (28). In contrast, highly intense powder X-ray diffraction patterns were obtained from the tapetum in regions surrounding the bottom parts, and immediately below the retina/rhabdom layer (red spot, Fig.3A). This confirms that the tapetum contains crystalline material and that the crystals have no preferred orientation. Raman spectra on thin cross-sections of eye tissue unambiguously identify the crystalline, birefringent material of the tapetal cells as almost pure isoxanthopterin (Fig. 3B). We also observed dark, absorbing granules lying below the tapetum which exihit a Raman signature characteristic of pigments (Fig. S3) (29).

**Fig. 3.**
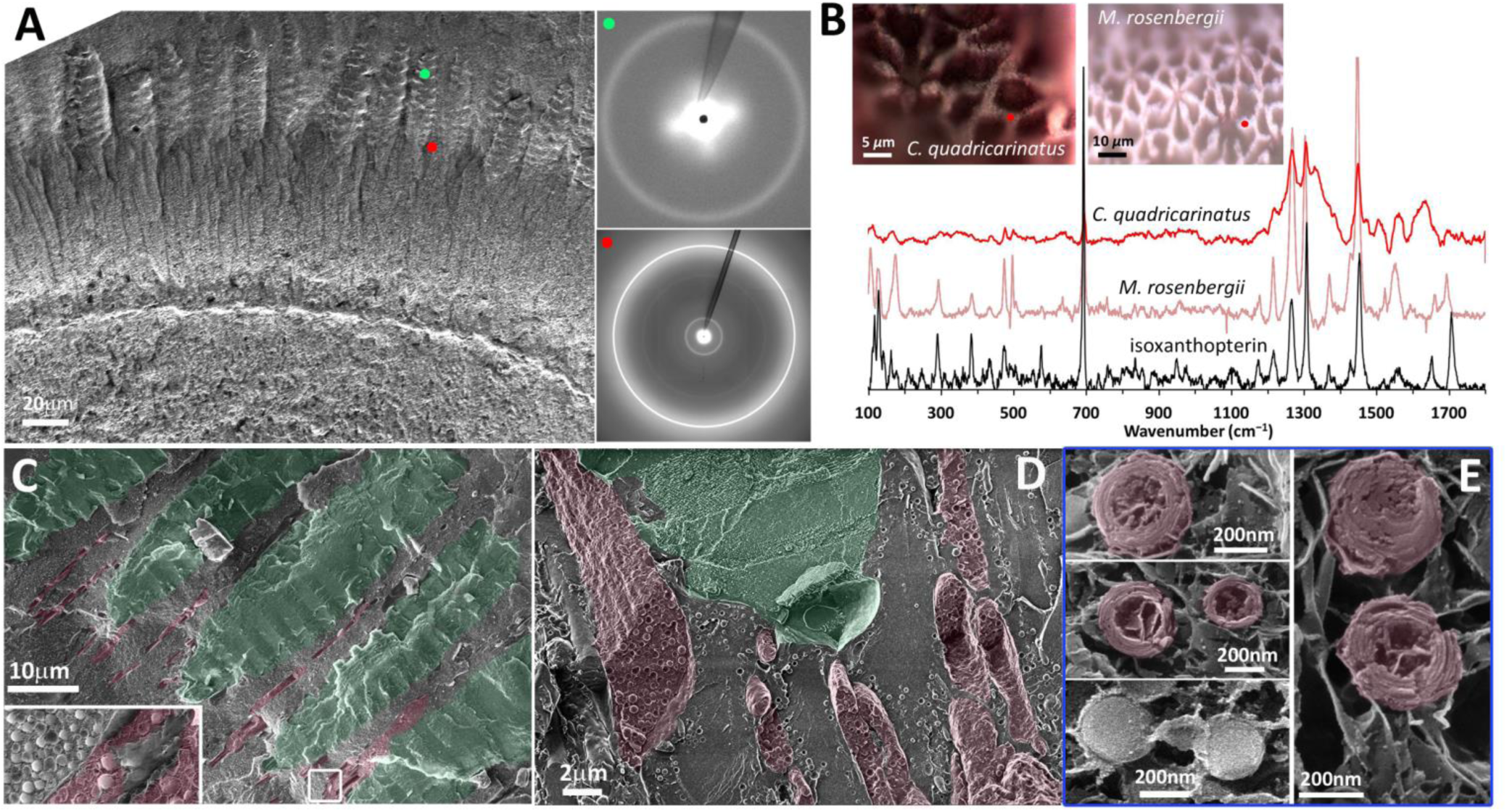
Physical, chemical and ultrastructural properties of the tapetum. (*A*) X-ray diffractograms (right) obtained from the rhabdom (green spot) and tapetum (red spot) using a micro-spot X-ray beam. The SEM image (left) shows the locations from which the diffractograms were obtained. (*B*) Raman spectra obtained from the white, birefringent tapetum material (red spectrum) of *C. quadricarinatus* and *M. rosenbergii* together with the spectrum of isoxanthopterin (solid black spectrum). Inset: polarized light microscope images of the tapeta, viewed from above, showing the locations (red spots) from which Raman spectra were taken. (*C*) – (*E*) cryoSEM images of cross-sections through the tapetum/rhabdom layer of *C. quadricarinatus*, viewed perpendicular to the eye axis. (C) Membrane bound extensions of the tapetal cells (pseudo-colored red) lying between the rhabdoms (pseudo-colored green). Inset: The tapetal cells are packed with nanoparticles [elucidated in more detail in (E)]. (*D*) The extensions of the tapetal cells, which contain a dense packing of nanoparticles, form a close-envelope around the rhabdom. (*E*) Fine structure of the crystalline tapetum nano-particles (red). Bottom left; in contrast, the absorbing pigment granules, found below the tapetum have a smooth internal texture.

To determine the ultra-structural arrangement of the tapetum at the nano-to microscale level, we performed cryo-SEM imaging on high-pressure-frozen, freeze-fractured eyes of *C. quadricarinatus* and *M. rosenbergii*. Cryo-SEM images of longitudinal eye-sections show membrane-bound extensions of the tapetal cells lining the lower parts of each rhabdom (Fig. 3C), matching the light microscopy observations of the birefringent tapetum material (Fig. S2). These elongated protrusions are typically 5 *μ*m wide and are densely packed with nano-particles with an average diameter of 390 nm (±33 nm, *N=* 44) (Fig. 3D, E). The outer parts of these hollow nano-particles are composed of small segments, which are arranged in an ‘onion-skin’ lamellae with an average shell thickness of 80 nm (±11.5 nm, *N*=35) (Fig. 2E). The same nanoparticles were also observed in *M. rosenbergii* (Fig. S4). These nano-particles appear to be the principal component of the tapetal cells, and based on spatially resolved X-ray and Raman measurements (Fig. 3A, B) we conclude that they are composed of crystalline isoxanthopterin. The tapetum nanoparticles are comparable in dimensions to the wavelengths of visible light and will therefore scatter light by Mie scattering. We calculated the ratio of back-to forward-scattered light using Mie theory for a 390 nm diameter nanoparticle, with a shell thickness of 80 nm, assuming an effective refractive index of *n* = 1.78 (*SI Methods* and Fig. S5). We find two peaks in the amount of back scattering of 10% and 25% at wavelengths of 470 nm and 700 nm.

**Fig. 4.**
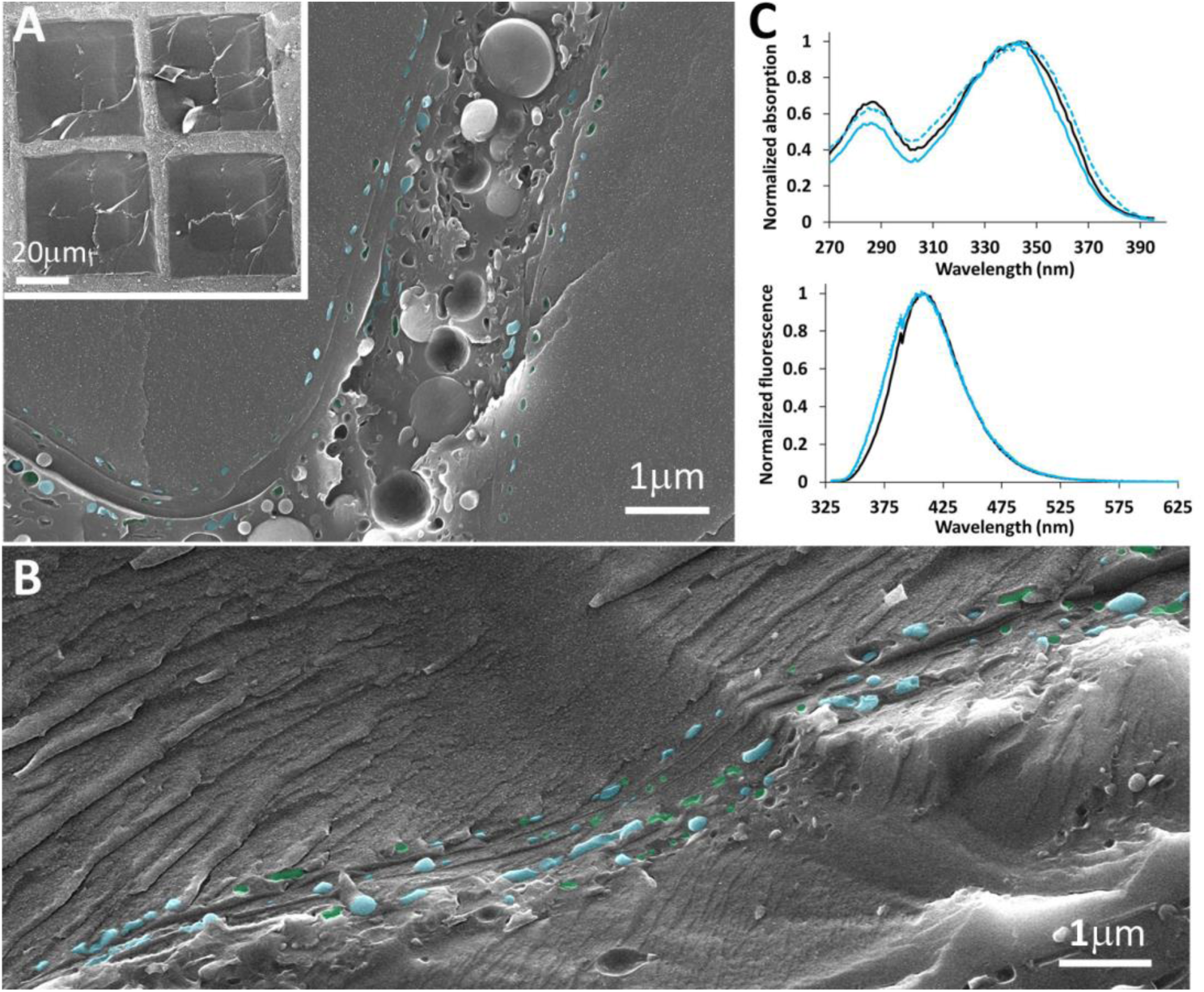
Ultrastructure and chemical properties of the multilayer mirror in *C. quadricarinatus* eyes. (*A*) Cryo-SEM image of the boundary between two ommatidia in the upper part of the eye viewed along the eye axis (inset; lower magnification cryoSEM image of the transverse section through 4 ommatidia). (*B*) Cryo-SEM image of the multilayer lining of an ommatidium from a longitudinal section through the eye. In (*A*) and (*B*) crystals are pseudo-colored blue and holes (produced by crystals being lost during freeze-fracture) are colored green. (*C*) UV-Vis absorption (upper) and emission (lower) spectra of the reflecting material extracted from the multilayer mirror (solid blue trace) and the tapetum (dotted blue trace) alongside the spectra obtained from isoxanthopterin crystals (black trace).

Cryo-SEM images in the area of the distal mirror in the upper part of the eye (Figs. 1C, 2A-C) reveal that each ommatidium is bound by a multilayer comprising 3 or 4 rows of solid objects interspersed with cytoplasm (Fig 4A, B). Polarized light microscopy strongly suggests that the objects are crystalline (Fig. 2C). Each row of the multilayer contains numerous crystals housed inside a membrane-delimited compartment surrounding the entire perimeter of an ommatidium. In the innermost rows, the crystals are plate-like (typically 200 nm wide and 40 nm thick) although there is no well-defined or conserved crystal morphology. The crystals are not closely packed and we estimate that in a single row there is about 50% coverage of crystals. A dense-packing of larger, *ca*. 1μm absorbing pigment granules (30, 31) occupies the central region between two ommatidia. These granules have a similar texture to the absorbing pigment granules found below the tapetum (Fig. 3E). The reflecting material extracted from both the multilayer and tapetum reflectors is highly fluorescent and exhibits excitation and emission peaks in the UV-Vis spectra characteristic of isoxanthopterin (22)(Fig. 4C). The simulated reflectance spectra derived from the crystal thicknesses and spacings measured by cryo-SEM (Fig. S6) show that the mirror exhibits broadband reflectivity and is 50–100% reflective for light impinging at angles of incidence between 0–30° (with respect to the ommatidial axis) but is only 20% reflective at normal incidence.

**Fig. 5.**
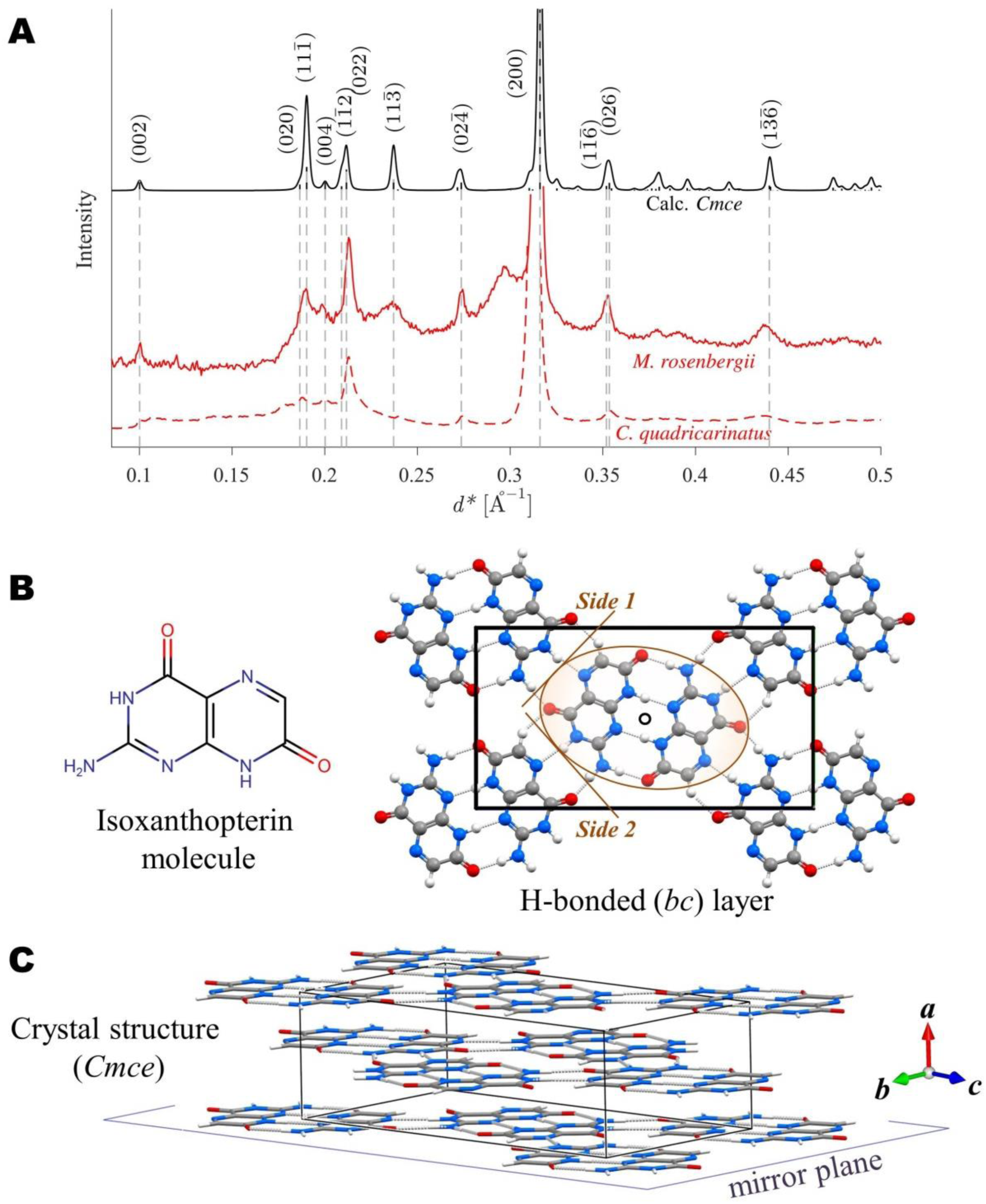
Crystal structure of isoxanthopterin. (*A*) X-ray powder diffraction profiles; calculated isoxanthopterin structure (black trace) alongside experimental data obtained from the tapetum of *M. rosenbergii* and *C. quadricarinatus*. The intensity of the strongest reflection (200) was truncated to make all the reflections more visible. Additional details in *SI text* and Fig. S8. (B) Molecular structure of isoxanthopterin and the crystal H-bonded layer. The brown contour indicates a centro-symmetric H-bonded dimer. (*C*) Calculated crystal structure of isoxanthopterin.

X-ray diffraction (XRD) patterns obtained from the crystals in *C. quadricarinatus* and *M. rosenbergii* tapeta exhibit sharp reflections indicative of well-ordered crystals (Fig. 5A). These experimental data form the basis for verifying the validity of the crystal structure of isoxanthopterin determined independently from crystal structure prediction calculations. To determine the crystal structure of isoxanthopterin, we utilized a similar approach to that reported previously for the crystal structure prediction of biogenic guanine (32). Both guanine and isoxanthopterin are planar conjugated molecules with numerous H-bond donor and acceptor sites. Our calculations on isoxanthopterin relied on the assumption that the crystal structure is dominated by intermolecular H-bonding and close-packing interactions. This assumption is supported by the presence of a very intense reflection with a *d*-spacing of ~3.2 Å in the X-ray diffraction patterns, suggestive of a closed-packed intermolecular layer arrangement. In the guanine XRD pattern, the 3.2 Å reflection corresponds to the distance between H-bonded layers of guanine molecules.

The first step in our crystal structure prediction was to construct a probable H-bonding motif for the isoxanthopterin molecules. The planar, molecule (Fig. 5B, left) contains four NH and one CH proton donors and six lone-pair electron lobes. We first form of a cyclic centrosymmetric dimer containing four H-bonds as the basic building block of the crystal structure (Fig. 5B, brown circle). We then take advantage of the H-bonding complementarity between the two adjacent sides of the isoxanthopterin molecule (Fig. 5B, right) to form a two-dimensional H-bonding array by twofold screw (2_1_) and (*c*) glide symmetry (Fig. 5B). By such means we generate a unique layer of H-bonded molecules of symmetry 2_1_/*c*. After generating different interlayer packing motifs, *via* monoclinic and orthorhombic symmetries we optimize these crystal structures using dispersion-inclusive density functional theory (DFT) *(SI methods)* and compare the calculated and observed XRD patterns.

The best fit between the observed and computed diffraction data was found for an orthorhombic structure, where the interlayer packing motif was generated from the planar H-bonded layer by an additional *b*-glide at *x=*¼ (parallel to the *bc* layer), yielding a new molecular layer at x=1/2, in the *Pbca* space group (Fig. S7). DFT-based geometry optimization of this structure yielded a higher symmetry structure, possessing the rarely observed space group *Cmce*, which corresponds to a perfectly planar H-bonded layer of isoxanthopterin molecules (Fig. 5C and *SI text*). The X-ray diffraction conditions for the *Cmce* space group fit well with the unusual experimentally observed XRD pattern of the biogenic samples, insofar as there is a region in reciprocal space (0.1<*d*^*^<0.2 Å^−1^) lacking any reflections (Fig.5A). Agreement between the computed and observed XRD data is excellent. A first principles calculation of the refractive index of isoxanthopterin, based on density functional perturbation theory, yields a high value of *n* = 1.96 for light incident along the *a* axis.

## Discussion

*In situ* chemical and structural analyses of the eyes of *C. quadricarinatus* and *M. rosenbergii* show unequivocally that the tapetum reflector is constructed from crystalline isoxanthopterin. UV-Vis analysis of the reflecting material extracted from the multilayer mirror of the upper distal reflector showed that this is also composed of isoxanthopterin and polarized light microscopy shows that the material is birefringent, and thus most probably crystalline. These results are consistent with earlier chemical analyses that isoxanthopterin is present in crustacean eyes (22). The crystal structure of isoxanthopterin is characterized by planar arrays of H-bonded isoxanthopterin molecules, reminiscent of the crystal structure of biogenic guanine. The extremely high calculated refractive index in the H-bonded plane, *n* = 1.96 makes isoxanthopterin an ideal reflecting material. In contrast to inorganic biominerals, very few types of functional organic crystals have been identified in animals. Crystalline guanine is found widely in the reflective systems of animals (2) and the purines xanthine and uric acid have been identified as the reflective materials in insect’s eyes (33) and cuticles (34) respectively. Pirie reported that the eye-shine of lemurs was produced by crystals of riboflavin (35). A key question is how many other reflecting organic crystals remain to be discovered in nature.

Isoxanthopterin belongs to the pteridine family of molecules, heterocyclic compounds composed of fused pyrimidine and pyrazine rings. Historically it was thought that the pteridines function exclusively as light-absorbing pigments intrinsically associated with yellow/red xanthophore pigment cells (36). Pteridine molecules are indeed used as pigments in a wide variety of animals and especially in insects, but not in a crystalline form. Here we present conclusive evidence of a crystalline pteridine in nature thus challenging the conventional functional distinction between the light-absorbing pteridines and light-reflecting (crystalline) purines. The discovery of crystalline isoxanthopterin in two distantly related decapod families (*C. quadricarinatus* and *M. rosenbergii* are from the *Astacidea* and *Caridea* infraorders respectively (37)) suggests that this material is widespread in the decapod crustaceans. There is indirect evidence that other animal tissues, including the avian iris (38) and the iridophores of amphibians (39) contain reflecting granules composed of pteridines.

## Tapetum

High-resolution cryo-SEM images of the eyes of *C. quadricarinatus* and *M. rosenbergii* show that the isoxanthopterin crystals of the tapetum are located inside *ca*. 400 nm nanoparticles comprised of concentric crystalline lamellae. Other ultrastructural studies found similar 400 nm “reflecting granules” in both the tapeta (25, 30, 31) and epidermis (40) of decapod crustaceans and Matsumoto (41) described pterinosomes (pteridine-containing granules) with a “concentric lamellar” structure in fish. It remains to be seen whether this texture, which seems to be a characteristic feature of pteridine granules in nature, is always indicative of the presence of crystalline particles (Fig. 3E). The tapetum nanoparticles we observed are comparable in dimensions to the wavelengths of visible light and will therefore scatter light by Mie scattering. The relatively high scattering efficiency of these particles is due to the high refractive index of isoxanthopterin crystals. Both *C. quadricarinatus* and *M. rosenbergii* live in turbid, shallow freshwater and approximately 10–12% of the yellow-green light which penetrates this environment will be back-scattered by an invidual nanoparticle (Fig. S5)(42). The scattering nanoparticles are densely packed inside ~5 *μ*m thick tapetal cell extensions that closely encapsulate the base of the rhabdom (Fig. 3C, D). Photons impinging on the tapetum will thus undergo multiple scattering events from numerous individual particles in these extensions resulting in a significant amount of light being back-scattered to the rhabdoms.

The principal function of the tapetum is to direct photons, which are not initially absorbed by the retina back to the retina, providing a second opportunity for photon capture. The ultrastructural arrangement of the tapetum, which forms a reflective sheath around the base of each rhabdom, also suggests a second function, namely to prevent optical cross-talk between rhabdoms (26, 27). In the dark-adapted reflection superposition eye, light is collected from a wide field of view. This could result in loss of resolution since light incident from the peripheral field of view, with steep angles of incidence might not remain within a single rhabdom. The protective sheath of the tapetum prevents such light being transmitted between adjacent rhabdoms. The tapetum thus serves to simultaneously increase the sensitivity and preserve the resolution of the eye. The fact that the tapetum sheath only partially encapsulates (25, 26) the rhabdom (Fig. 3C) may also be functional, since a fully encapsulated rhabdom may prevent sufficient light from reaching the target, lowering eyes sensitivity (26). In light-adapted eyes, proximal absorbing pigments, housed within the retinal cells migrate from below the basal membrane and completely envelope the tapetum sheath (25, 43) preventing the rhabdoms from being damaged by over-exposure to photons (43, 44).

Diffusely scattering tapeta have also been observed in fish (45), crocodiles (46) and in the opossum (47), and are constructed from a range of crystalline and non-crystalline materials (48). The light scattering sub-micron sized guanine-crystals found in the eyes of the elephant nose fish (49) increase the amount of light absorbed by co-localized melanin pigment granules, enabling simultaneous activation of both rod and cone cells in the same ambient light conditions. Wilts *et al* reported (50) that the unusually bright colors of *Pieridae* butterfly wings are produced by efficient Mie-scattering from ellipsoid granules of pteridines with an extremely high refractive index. They conclude that the high refractive index is due to a high-density of pteridines which they assume must approach a “close-packing” assembly in the granules.

## Distal mirror

The multi-layer reflector in the upper part of *C. quadricarinatus* and *M. rosenbergii* eyes is formed from 3 or 4 rows of sparsely populated plate-like bodies which we assume are nano-crystals of isoxanthopterin. Since grazing incidence light (< ~15°) is reflected from the walls of the ommatidia by total internal reflection (15, 17), the role of the distal mirror is to reflect light entering at higher incidence angles, effectively increasing the pupil size and sensitivity of the dark-adapted eye. Reflectivity simulations assuming their crystalline nature show that the mirror is 45–70% reflective for light entering the ommatida at 15–30° but drops to about 20% for light impinging at normal incidence due to the low packing-density of the crystals and the small number of rows in the multilayer. The pigments lying between ommatidia will absorb any light not reflected by the mirror at such angles. We note that our calculation of broadband reflection in the *C. quadricarinatus* mirror is inconsistent with Land’s observations of the distal mirror in an oceanic shrimp. Land (15) observed that it had a “green-specular” appearance when viewed at near normal incidence. This difference might be accounted for by the different light conditions in the habitats of these two species.

We suggest that the poor performance of the mirror at very high angles of incidence may actually be functional in the dark-adapted eye. Light entering the eye at extreme angles of incidence will be particularly affected by spherical aberrations and as such will contribute disproportionally to image blurring (27, 51). A mirror that becomes progressively less efficient at very high angles will contribute less light from the peripheral field of view, thus mitigating against image blurring – a process known as apodization in optics. The poor performance of the mirror at high angles would also prevent light undergoing multiple reflections in a single ommatidium, which would lead to light being scattered stochastically across the clear zone onto different parts of the retina. Much like the tapetum then, the ultrastructural organization of the distal mirror seems to be tuned to optimize the balance between increasing the sensitivity of the eye on the one hand and maintaining resolution on the other(26, 27).

## Conclusions

Surprisingly little is known about functional organic crystals in nature and few of these materials have been identified. Our discovery that crystalline isoxanthopterin is the highly reflective optical material in the eyes of decapod crustaceans paves the way for further research in the emerging field of “organic biomineralization” and suggests that other such materials, together with their optical properties may remain to be discovered. Our results also call for pteridines found more widely in the animal kingdom to be investigated and their possible role as biological reflectors to be re-examined. Eyes based on reflective optics are often extremely light-sensitive and are found in animals inhabiting environments where light is at a premium. However, in such wide-aperture eyes there is often a trade-off between sensitivity (e.g. increasing the field of view) and resolution (e.g. spherical aberration). The ultrastructural organization of the reflectors in the dark-adapted eyes of decapod crustaceans appear to be tuned to optimize the balance between increasing light sensitivity and maintaining image resolution.

## Acknowledgements

We thank Prof. Ashwin Ramasubramaniam for his helpful advice on calculating the refractive index of isoxanthopterin and Neta Varsano, Nadav Elad and Batel Rephael for help with related experiments. Electron microscopy studies were supported by the Irving and Cherna Moskowitz Center for Nano and Bio-Nano Imaging at the Weizmann Institute of Science. This work was supported by the Israel Science Foundation (Grant 583/17) and the Crown Center of Photonics and the ICORE: The Israeli Excellence Center “Circle of Light”. B.A.P. is the recipient of a Human Frontiers Science Program - Cross-Disciplinary Postdoctoral Fellowship. L.A. and S.W. are the incumbents of the Dorothy and Patrick Gorman Professorial Chair of Biological Ultrastructure and the Dr. Trude Burchardt Professorial Chair of Structural Biology, respectively.

